# Primate dopamine neurons activated in rewarding environment by tonic disinhibition from amygdala

**DOI:** 10.1101/2022.03.28.486109

**Authors:** Kazutaka Maeda, Ken-ichi Inoue, Masahiko Takada, Okihide Hikosaka

## Abstract

Seeking out good and avoiding bad objects is critical for survival. In practice, objects are rarely good every time or everywhere, but only at the right time or place. Whereas the basal ganglia (BG) are known to mediate goal-directed behavior, for example, saccades to rewarding objects, it remains unclear how such simple behaviors are rendered contingent on higher-order factors, including environmental context. Here we show that amygdala neurons are sensitive to environments and regulate dopamine (DA) neurons via an inhibitory projection to substantia nigra pars reticulata (SNr) neurons. We combined optogenetics and multi-channel recording to demonstrate that rewarding environments induce tonic firing changes in DA neurons. These tonic responses are mediated by disinhibition via GABAergic projection onto DA neurons from SNr, which in turn are suppressed by inhibitory projection from the amygdala. Thus, the amygdala provides an additional source of learning to BG circuits, namely contingencies imposed by the environment.

## Introduction

We choose actions in accordance with our goals, for instance to seek out good (rewarding or useful) objects and avoid bad (harmful or useless) objects. Goal-directed actions commonly begin with eye movements towards good objects or away from bad objects. How is such goal-directed behavior controlled by the brain? Eye movements associated with approach and avoidance are controlled by two parallel pathways in basal ganglia (BG) circuits (Amita et al., 2019; Hikosaka et al., 2019a). Approach behavior is mediated by the direct pathway from the caudate nucleus (CD) to the superior colliculus (SC) via the substantia nigra pars reticulata (SNr), whereas avoidance behavior is mediated by the indirect pathway through the globus pallidus externus (GPe), i.e., the CD-GPe-SNr-SC pathway. These direct and indirect pathways originate from the caudate head (CDh) and the caudate tail (CDt), which process short- or long–term memory, respectively (Kim et al., 2015; Hikosaka et al., 2019b).

In real life, different objects are typically found in different environments. Thus, the act of encountering a new environment itself conveys predictive information about what objects are likely to be found therein and accordingly influences behavior (Maren et al., 2013; Maeda et al., 2018; Kunimatsu et al., 2021). How does the brain process such predictive environmental information? We hypothesized that brain regions are likely to play roles that both encode distinct environments and influence the BG circuits. The amygdala is a key structure for encoding emotionally significant environments (Murray, 2007; Choi and Kim, 2010; Tamietto and de Gelder, 2010; Amir et al., 2015; Saez et al., 2015; Tovote et al., 2015; Maeda et al., 2018; Maeda et al., 2020). Furthermore, the amygdala sends output to BG, including CDt, GPe, and SNr (Price and Amaral, 1981; Shinonaga et al., 1992; Vankova et al., 1992; Fudge and Haber, 2000; Griggs et al., 2017). In the present study, we investigated what information is conveyed through the amygdala-nigra projection using multi-channel recording probes combined with pathway-selective optogenetic technique.

## Results

### Dopamine activity for environmental contexts

Pavlovian conditioning procedures that pair conditioned stimuli (CS) with unconditioned stimuli (US) have widely been used for testing dopamine (DA) neurons (Schultz, 2016). We designed a Pavlovian procedure in which each trial started with the onset of a distinct environment that constrained what subsequent CS objects and US events could occur. In possible reward (PR) environments, the US took the form of a liquid reward, the occurrence of which was not guaranteed (Fig. 1a). By contrast, no reward (NR) environments signaled the possibility of an airpuff (Fig. 1b). In both environments, the CS were represented by unique fractal objects associated with their corresponding US with 100%, 50%, or 0%. Fractal CS objects appeared 1.8 s after environment onset. After 1s, the US occurred (or were omitted) and simultaneously the CS and the environment background image disappeared.

**Fig.1.**
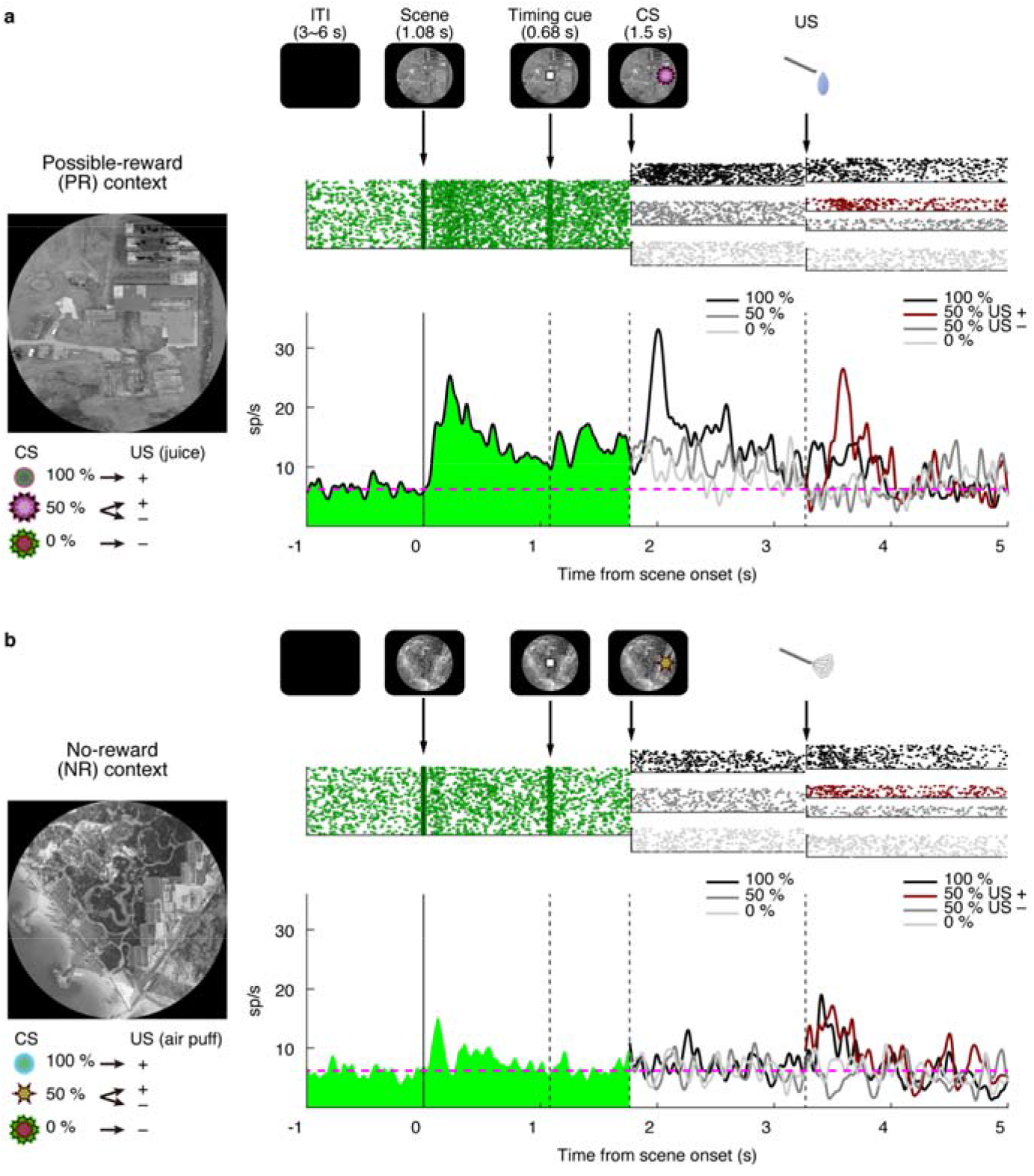
DA response to environmental contexts. **a**, Responses of a single DA neuron in the PR environment of Pavlovian task with scene. In the PR environment, a particular scene was presented, and three conditioned stimuli were associated with water with 100%, 50% and 0% probability, respectively. **b**, Responses of the same DA neuron in the NR environment of Pavlovian task with scene. In the NR environment, another scene was presented, and other three conditioned stimuli were associated with air-puff with 100%, 50% and 0% probability, respectively. The environments were pseudo-randomly switched within the block. The trial started after the presentation of a scene image (diameter: 50 degrees) on the screen. After 1.08 s, a timing cue was presented on the scene. After 0.68 s, the timing cue disappeared, and one of the three conditioned stimuli was presented in each environment. After 1.5 s, the conditioned stimulus disappeared, and the unconditioned stimulus (water reward in PR or airpuff in NR) was delivered. Actual scenes and fractals for CSs used are shown on the left. The top row shows a raster plot of firing for the scene and each CS and US in the set, with dots indicating spike times. The bottom row shows the average firing rate in the set. ITI, inter-trial interval; PR, possible-reward; NR, no-reward; CS, conditioned stimulus; US, unconditioned stimulus; DA, Dopamine.

After the monkey learned the task, we recorded single-unit activity from the substantia nigra pars compacta (SNc). When the PR-environment appeared, DA neurons showed a phasic response followed by tonic excitation (Fig. 1a). On the other hand, no tonic response was observed in the NR-environment, but DA neurons responded to CS and US phasically (Fig. 1b). In the PR-environment, the response to CS was largest when reward prediction was largest (100%), but the response to US was largest when reward outcome was uncertain (50 %). This pattern is typical of DA responses to CS in standard Pavlovian tasks, that is, the absence in the environmental context (Schultz et al., 1997; Matsumoto and Hikosaka, 2009). These data suggest that DA neurons integrate two kinds of information: Environment in the tonic response and Object in the phasic response.

### Amygdala-nigra projection and DA neurons

We hypothesized that the amygdala is a source of the information about environment. Recent studies showed that amygdala neurons send their axons to SNr and modulate neuronal responses therein (Maeda et al., 2020; Steinberg et al., 2020). To study whether DA neurons also receive amygdala input, we injected AAV vector (AAV2-CMV-ChR2-EYFP) into the amygdala, mostly in the central nucleus (CeA) (Fig. 2a) to drive ChR2 expression in amygdala neurons and their axons terminating within the nigra (Fig. 2b). If DA neurons receive input from the amygdala, optical stimulation in the nigra would inhibit or excite DA neurons. For both electrophysiological recording and optical stimulation, we used a multi-site linear electrode with optic fiber ports inserted in either the amygdala (Fig. 2a) or the nigra (Fig. 2b). The nigra array had three optical stimulation sites, permitting optical stimulation at variable distances from the recording contacts (Fig. 2b). Then, we recorded the activity of many neurons while optically stimulating (duration: 200 ms) at one position in each trial. The stimulation intensity was set low enough to avoid irrelevant effects by optical stimulation (see Methods and Fig. S1). For the amygdala electrode, the effect of optical stimulation would mostly act on the cell somata of amygdala neurons, whereas for the electrode in the nigra, the effect of optical stimulation would excite the axon terminals of amygdala neurons.

**Fig. 2.**
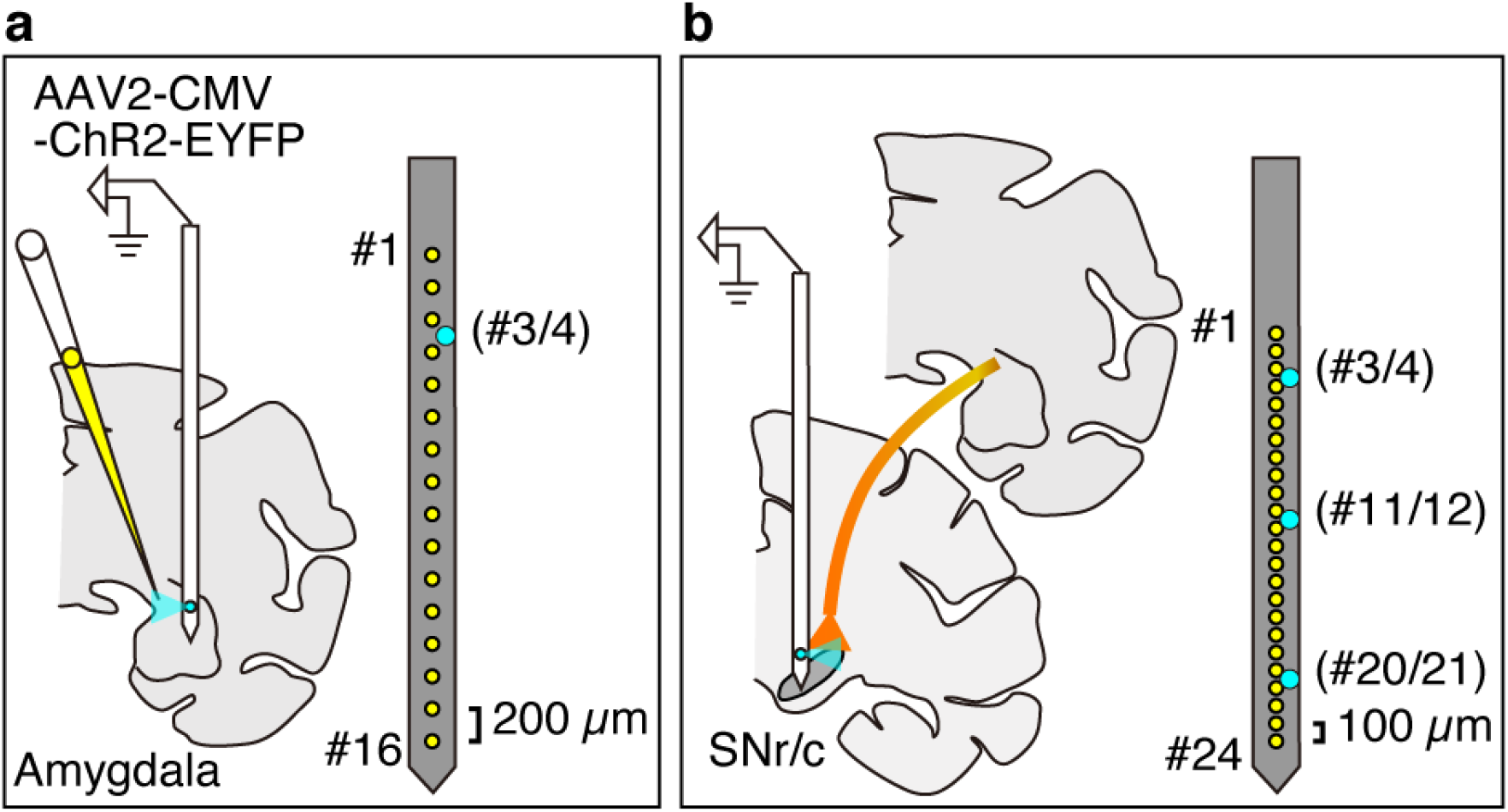
Optogenetic stimulation of amygdala-nigra pathway. **a**, Injection of the viral vector (AAV2-CMV-ChR2-EYFP) into the amygdala and stimulation/recording of amygdala neurons via a 16-contact probe (yellow dots, 200 µm interelectrode spacing) with a fiber optic which is placed between contact 3 and 4 (cyan dot). **b**, Stimulation of ChR2 expressing amygdala axon terminals in the substantia nigra via a 24-contact probe (yellow dots 100 µm interelectrode spacing) with three fiber optics (placed between contacts 3 and 4, contact 11 and 12, and contact 20 and 21; cyan dots).

Many amygdala neurons were modulated by optical stimulation in the amygdala (n = 76/211, 36%), mostly with excitatory responses (n = 53/76, 70%, Fig. 3a-left, Fig. 3b: red dots), confirming that excitatory opsin was successfully expressed locally. Some neurons were inhibited (n = 23/76, 30%, Fig. 3a-right, Fig. 3b: blue dots), which might not be due to the direct effect of optical stimulation on these cells, but was presumably driven by local inhibitory network of ChR2-expressing neurons. Consistent with this interpretation, inhibited neurons were less tightly clustered around the stimulation site than excited neurons (Fig. S2d), and inhibition occurred at longer latency than excitation (Fig. 3a-right).

**Fig. 3.**
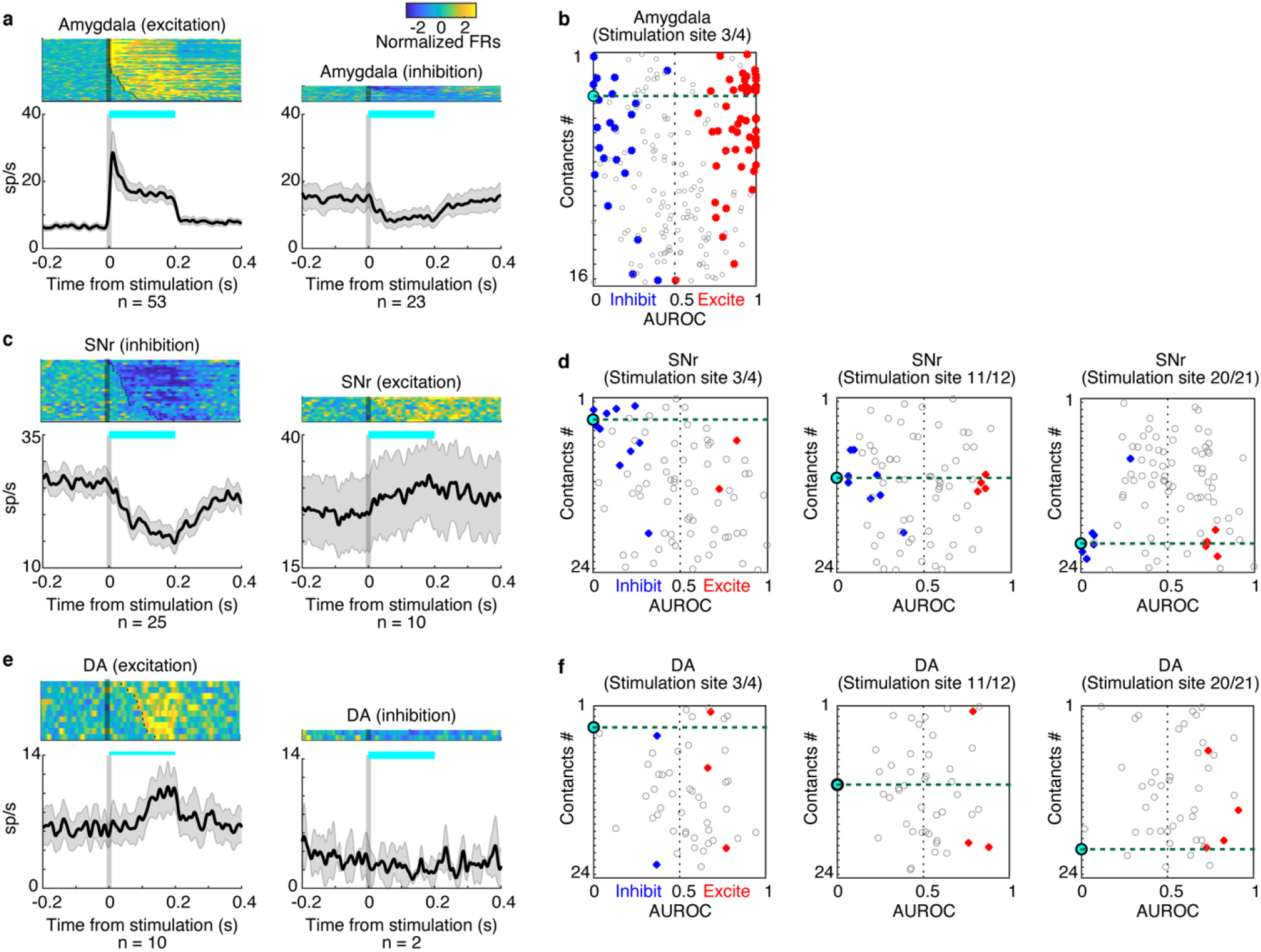
Neuronal modulation by optogenetic stimulation. **a**, Population activity of the amygdala neurons that were excited (left) or inhibited (right) by optogenetic stimulation. Spike activity was smoothed with a Gaussian kernel (σ = 10 ms). The shaded gray area indicates ± 1 SEM for the averaged population activity. Responses of individual neurons (upper) were converted to color scale and sorted by modulation latencies (dots). The duration of optical stimulation was 200 ms (cyan bar). **b**, Data from combined all sessions recording amygdala neurons while optogenetically stimulating the amygdala. The degree of the modulation by the stimulation is defined as the area under the receiver operating characteristic curve (AUROC) based on the activity in stimulation trials vs. control (sham stimulation) trials for each neuron. Each dot indicates the AUROC score in each neuron. Red or blue color shows statistically significant changes of the neuronal responses by optogenetic stimulation (P < 0.05, t-test). AUROC >0.5 or <0.5 indicates that the neuron was excited or inhibited by stimulation comparing with non-stimulation trials, respectively. **c, e**, Population activities of SNr or DA neurons that were inhibited or excited by optogenetic stimulation of amygdala’s axon in substantia nigra. **d, f**, Data from combined all sessions recording SNr or DA neurons while stimulating the amygdala’s axon in the substantia nigra by one of stimulation ports of fiber optic (left: 3^rd^/4^th^, center: 11^th^/12^th^, right: 20^th^/21^st^). The same formats of d.

Next, we optically stimulated amygdala axons in the nigra while recording SNr and DA neurons (Fig. 2b). In terms of the characteristic of DA neurons, we measured their spike shape and compared with that of SNr neurons. The data showed that the spike shape of DA neurons was significantly wider than that of SNr neurons (Fig. S3). This is consistent with previous data (Matsumoto and Hikosaka, 2009). Most of the affected SNr neurons exhibited inhibitory responses (n = 25/35, 71%, Fig.2e-left), attributable to monosynaptic projections from GABAergic CeA neurons (Tepper et al., 1995). Inhibited neurons tended to be found on channels closest to the stimulation port (Fig. 3c-left, Fig. 3d, blue dots). Switching to a different stimulation port led to responses in a new population of nearby neurons (Fig. 3d, green dot line). Optical stimulation drove excited responses in a minority of SNr neurons (10/35, 29%, Fig. 3c-right, Fig. 3d, red dots), which tended to be weaker in magnitude (Fig. S2b). This pattern of responses could be explained either by a small number of excitatory projections from the amygdala (areas adjacent to CeA), or by disinhibition resulting from disynaptic activation of SNr neurons. In contrast to SNr neurons, most DA neurons were excited by optical stimulation (10/12, 83%, Fig. 3e-left), and responsive channels were observed irrespective of the proximity to stimulation ports (Fig. 3f, red dots). This spatial pattern suggests that the effect of optical stimulation on each SNr neuron is highly localized to the axon terminals of amygdala neurons that have synaptic inputs to the SNr neuron. By contrast, the effect of optical stimulation on each DA neuron is spatially variable because the disynaptic inputs from SNr neurons depend on the spatial variation of their axons.

Further insight into circuit organization can be gleaned by assessing the latency of neuronal responses evoked by optical stimulation. SNr neurons were activated at shorter latency than DA neurons (Fig. 4a), again suggesting that SNr neurons are the primary target of optical stimulation. Moreover, latency varied systematically as a function of the distance from the stimulation port for SNr neurons (Fig. 4e), but not for DA neurons (Fig. 4f). Among optically activated SNr neurons (Fig. 4e), cells that were close to the optical stimulation (within 0.25 mm; n=17/25, 68 %) were inhibited quickly (mean: 47.6 +- 31.9 ms). The remaining neurons beyond 0.25 mm (n=8/25, 32 %) were inhibited significantly, but more slowly (mean: 93 +- 51.3 ms, t-test _P_ = 0.0453).

**Fig. 4.**
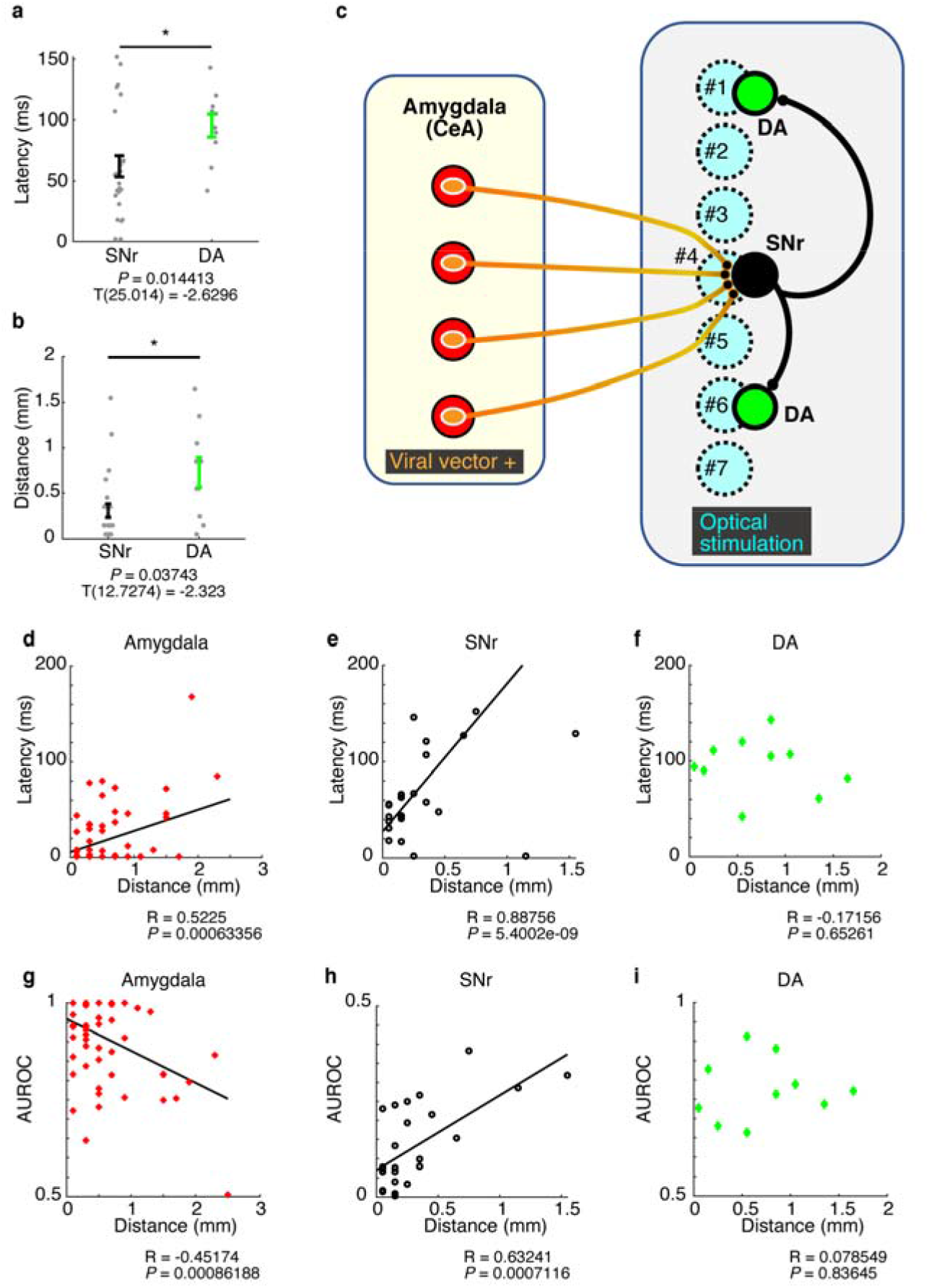
Amygdala-SNr-DA circuit and environmental information. **a**, Averaged latency for the neuronal modulation started by optogenetic stimulation. Error bars show ± 1 SEM, and each gray dot represents one neuron. Responses are inhibited for SNr neurons and excited for DA neurons. **b**, Distance between effective optical stimulation and recording sites. Error bars show ± 1 SEM, and each gray dot represents one neuron. **c**, Schematic showing effect of optogenetic stimulation on the amygdala - substantia nigra circuit. **d-f**, Relationship between distance (abscissa) and latency (ordinate) in each area. Excited neurons in the amygdala, inhibited neurons in SNr, and excited neurons in DA are shown in d, e, and f, respectively. **g-i**, Relationship between distance (abscissa) AUROC (ordinate) in each area. AUROC close to 1 means stronger excitation, while AUROC close to 0 means stronger inhibition (compared with AUROC 0.5). Same population as d-f.

These data suggest the direct connections of amygdala neurons to SNr neurons, as shown in Fig. 4c. In this schematic diagram, four amygdala neurons (shown by red circles) are infected with viral vector (orange ovals) and express ChR2 in their axons (orange lines). If these cells project to a single SNr neuron (black circle), their axons are close with each other to make synapses to the SNr neuron. Then, if optical stimulation is delivered at a short distance from the synapses, most axons (e.g., the four shown near cyan circle #4) will be activated and cause a strong and short-latency inhibitory postsynaptic potential (IPSP) in the SNr neuron. Alternatively, if the optical stimulation site is remote from the SNr neuron (cyan circle #1, 2, 3, 6, or 7), no axon to the SNr neuron will be activated. If the axonal bundle converging on the SNr neuron is only partially activated (e.g., the one fiber at cyan circle #5), a weaker and longer-latency IPSP will be evoked (Fig. 4e, Fig. 4h).

In contrast, neither the latency (Fig. 4f) nor the magnitude (Fig. 4i) of DA neuron responses showed any relation to the distance between optical stimulation and recording sites. These data suggest that DA neurons rarely receive direct inputs from the amygdala. Thus, optical stimulation close to a DA neuron (Fig. 4c, cyan circle #1 or 6) will not activate or inhibit the DA neuron because the incoming axons do not express ChR2 transported from the amygdala. Instead, if DA neurons receive inputs from an SNr neuron (as shown in Fig. 4c), the optical stimulation close to the SNr neuron (cyan circle #4 or #5) may cause synaptic effects on the DA neurons (if the SNr neuron has synaptic connections to the DA neuron). Accordingly, the optical stimulation would affect DA neurons that are located at random positions depending on the SNr-DA circuits (as illustrated in Fig. 4c). As a whole, the effect of the optical stimulation on DA neurons would therefore have no relation between distance and latency (Fig. 4f) nor between distance and activity (Fig. 4i).

Taken together, these data shed light on the organization of the amygdala-SNr-DA neuron circuit. Our results indicate that both the amygdala-SNr and the SNr-DA neuron connections are basically inhibitory. Thus, the main influence of the amygdala on DA neurons is excitatory and mediated by disinhibition through SNr neurons (Figure 2c, e, g; left side). In fact, anatomical studies have shown that the main projection from the amygdala to the nigra is inhibitory and originates from CeA (McDonald, 1982; Pitkanen and Amaral, 1994; Tovote et al., 2015). Likewise, neurons in SNr are well known to be inhibitory (Oertel et al., 1982; Parent et al., 1983; Hikosaka and Wurtz, 1985; Haber and Fudge, 1997) and project to DA neurons (Tepper et al., 1995; Kim et al., 2015), in addition to SC neurons (Hikosaka and Wurtz, 1983).

### Responses of amygdala, SNr, and DA neurons to environmental contexts

Thus far we have seen that DA neurons are sensitive to environmental contexts in addition to objects (Fig. 1). Optogenetic probing of functional circuitry then raised the possibility that the amygdala provides environmental information to DA neurons (Figs 2-3). In light of this hypothesis, we proceed here to evaluate the impact of environmental contexts on amygdala, SNr, and DA neurons in the Pavlovian task.

Many neurons showed tonic firing rate changes following the onset of PR-environment, to varying degree across the three areas (Fig. 5, S4). The amygdala neurons displayed tonic excitation (n = 62, Fig. 5a) more often than tonic inhibition (n = 8, Fig. S4a-left, also Fig. 5b). These neurons were recorded mostly in the dorsal part in the region near CeA (Fig. 5c-right). The SNr neurons exhibited tonic inhibition (n = 17, Fig. 5d) more often than tonic excitation (n = 7, Fig. S4b-left, also Fig. 5e), whereas vice versa for the DA neurons (tonic excitation: n = 32, Fig. 5g; tonic inhibition: n = 9, Fig. S4c-left, also Fig. 5h). The DA neurons with tonic excitation were found relatively dorsal part of the nigra (Fig. 5i).

**Fig. 5.**
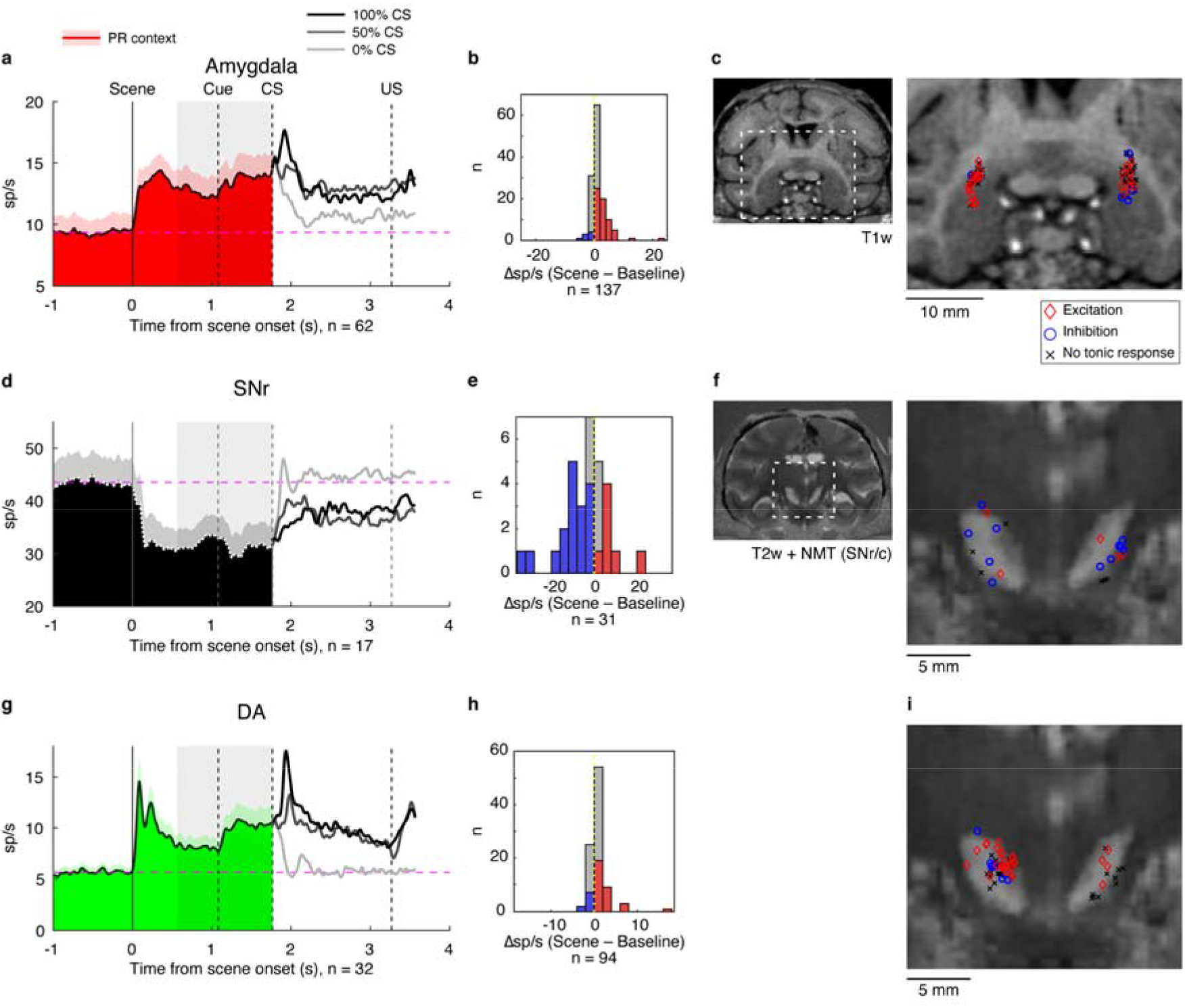
Tonic modulation in environmental contexts. **a, d, g**, Population activity for amygdala (excited responses), SNr (inhibited responses), and DA (excited responses) in the Pavlovian task (PR environment). The shaded gray area indicates spike-count window used to calculate average activities for the tonic component of the scene responses (0 – 1.2 s before the CS onset). Tonic excitation was shown in many of amygdala and DA neurons. Tonic inhibition was shown in the many of SNr neurons. The shaded red area indicates + 1 SEM for the averaged population activity. b, e, h, Distribution of responses to PR scene for amygdala, SNr, and DA neurons of all types (excited, inhibited, and non significant. The difference in firing rates was calculated by subtracting the baseline response (500 ms before scene onset) from the tonic response in each neuron. **c**, Recording locations of amygdala neurons in Monkey SO superimposed on MRI image. Dotted region shows location of magnified detail. Recording sites showing excited and inhibited responses to PR scene are indicated by red and blue markers, respectively (P < 0.05, T-test). Cells with no significant response to PR are indicated by black markers. **f, i**, Recording locations of SNr or DA neurons in Monkey SO. The same format as in c.

The overall results suggest that amygdala neurons may relay information about the context to the nigra, which in turn may cause these downstream neurons to distinguish between PR/NR scenes and associated CS fractals. To test this idea explicitly, we compared neuronal activities with the effects of optical stimulation which were recorded during the Pavlovian task. In two out of three cases, SNr neurons that were tonically inhibited by the environment were also inhibited by the optical stimulation. Likewise, three out of eight DA neurons tested had congruent excitatory responses both during the task and to the optical stimulation (Fig. S5). These data further support the interpretation that the information about environmental contexts is sent from the amygdala to DA neurons through disinhibition via SNr neurons.

Notably, the tonic firing rate changes induced by PR environments in all three areas (both excitatory and inhibitory responses) persisted as long as the possibility of reward remained. However, this persistent activity was truncated by the onset of the 0% CS that foreclosed the possibility of reward on the current trial (Fig. 5; Fig. S4). This finding indicates that tonic responses to the environment were not simple visual responses, but were rather associated with motivation (i.e., expectation of reward outcome).

We further considered whether such different neuronal responses to PR and NR scenes could be attributed to visual stimulus features rather than value information. To test this notion, the Pavlovian task was repeated with a second set of images for the PR condition, and the pattern of responses was compared across the two sets (Fig. S6a). The responses to the two PR-environments were significantly correlated in these sets for the amygdala, SNr, and DA neurons (P < 0.05, Pearson’s correlation, Fig. S6b-d). Our results confirm that these neurons were indeed sensitive to environmental information, rather than the visual features of individual scene images. These data further indicate that a subset of DA neurons mainly showed tonic excitation in the appetitive PR-environment like amygdala neurons, forming a striking contrast to tonic inhibition of SNr neurons, which is in harmony with the pattern of responses to the optogenetic stimulation (Figs. 3 and 4).

## Discussion

We used optogenetics and multi-channel electrophysiological recording to reveal that amygdala, SNr, and DA neurons are sensitive to environmental context. Specifically, we have found that tonic activity changes in DA neurons are generated by the amygdala-derived indirect input through SNr. These data suggest that the amygdala regulates DA cell activity in order to maintain motivational conditions in particular environments.

### Brain areas controlling BG circuits

Previously, it has been shown that BG control choices of objects for the sake of reaching a desired outcome (e.g., reward). This function is controlled by two sets of parallel BG circuits, namely the CDh circuit for flexible value and short-term memory, and the CDt circuit for stable value and long-term memory (Kim and Hikosaka, 2013). For both circuits, the direct pathway mediates the choice of good objects, whereas the indirect pathway mediates the rejection of bad objects (Amita and Hikosaka, 2019). These parallel pathways are necessary to make appropriate decisions, for instance, by taking advantage of the past experience with objects when possible. However, the goals of behavior can be contingent on diverse information in addition to a particular targeted object, such as the animal’s current environment. Optimal behavior requires that brain areas encoding such higher-order factors connect to decision-making areas that integrate information from multiple sources and ultimately guide behavior. Our results suggest that the amygdala encodes environmental information and sends it to the BG circuits to modulate SNr and DA neuron responses. Consistent with this idea, previous studies suggest that the amygdala may encode environmental information related to emotion (e.g., “wonderful”, “worried”, etc. (Everitt et al., 1991).

### Why Environment is important?

Animals often choose or reject particular objects to achieve favorable outcomes (e.g., reward). Previously the mechanisms of such goal-directed behavior have mostly been studied using very simple laboratory tasks. However, good or bad objects are often not easy to find in real life and may be located only in particular environments. For example, many animals respond to changes in food availability by seeking out new environments (Bauer and Hoye, 2014; Riotte-Lambert and Matthiopoulos, 2020). While favorable environments do not guarantee favorable outcomes, they often increase the probability of findings sought after objects or resources of interest. Accordingly, entering a noteworthy environment can prime specific goal-directed behaviors and neural pathways even before animals encounter objects for the goal. In this study, we used large visual images to represent environments that predict different emotional outcomes (e.g., reward). Could the persistent tonic activity change we observed be attributed to the persistence of one large visual image for the duration of each trial? Two findings argue against this interpretation. First, tonic activity was halted by the onset of the 0% CS fractal that foreclosed the possibility of reward, despite the continued presence of the large background image (Fig. 5, S4). Second, the same tonic activity changes were induced by two PR images with very distinct visual features (Fig. S6). Thus, SNr and DA neurons only responded to environmental stimulus images insofar as they predicted likely outcomes.

### Environment encoded by DA neurons

Most previous physiological investigations on DA neuron activity focused on their phasic components, for instance, responses to reward outcomes or reward-predicting visual stimuli (Schultz et al., 1997; Matsumoto and Hikosaka, 2009). Here we have found that DA neurons can also respond to stimuli with tonic activity changes (see Figs. 1 and 5). Likewise, studies in rodents reported the tonic modulation of DA concentration measured by microdialysis or voltammetric methods in various environments (Howe et al., 2013; Collins et al., 2016; Hamid et al., 2016). In these experiments, rodents locomoted through spatially expansive environments, in a manner analogous to the large visual stimuli used as environments in the current study. This suggests that entering a new environment can induce tonic changes in DA neurons in many animals, including at least both monkeys and rodents. Other studies in monkeys also demonstrated the tonic modulation of DA neuron responses which occurred under uncertain value conditions or during continuous changes of expected reward outcome (Fiorillo et al., 2003; Bromberg-Martin et al., 2010; Kobayashi and Schultz, 2014; Wang et al., 2021).

Environmental and various underlying contexts (e.g., uncertainty or continual change) are important in real life for all animals (Sih et al., 2011; Owen et al., 2017). Moreover, environmental information is related to the therapeutic effectiveness for some diseases, including Parkinson’s disease (Lim et al., 2005; Kwakkel et al., 2007). It is suggested that animals exposed to enriched environments exhibit resistance to parkinsonian symptoms (Nithianantharajah and Hannan, 2006). Furthermore, environments affect the sensitivity to some drugs that are believed to target midbrain DA neurons (Siegel et al., 1982). Our findings suggest that environmental information could promote neuronal activation, signaling, and plasticity through DA circuits, with important consequences for human and animal wellbeing.

### Mechanisms of amygdala-SNr-DA circuit

We propose that the amygdala operates on two distinct circuits within BG, with functional consequences manifest on both an immediate and a long-term time scale. Behavior executed over the course of a single trial is achieved on the immediate time scale and mediated by the pathway representing “Action in the Environment” in Fig. 6. We previously showed that amygdala neurons respond to emotional environments tonically and control eye movements via a direct inhibitory pathway to SNr (Maeda et al., 2018; Maeda et al., 2020). Saccades are triggered through disinhibition when the tonic inhibitory influence of SNr on SC is paused (Hikosaka and Wurtz, 1985). However, this mechanism alone is insufficient to account for complex behavior, such as learning from the past experience, or for taking uncertainty or environmental contingencies into account.

**Fig. 6.**
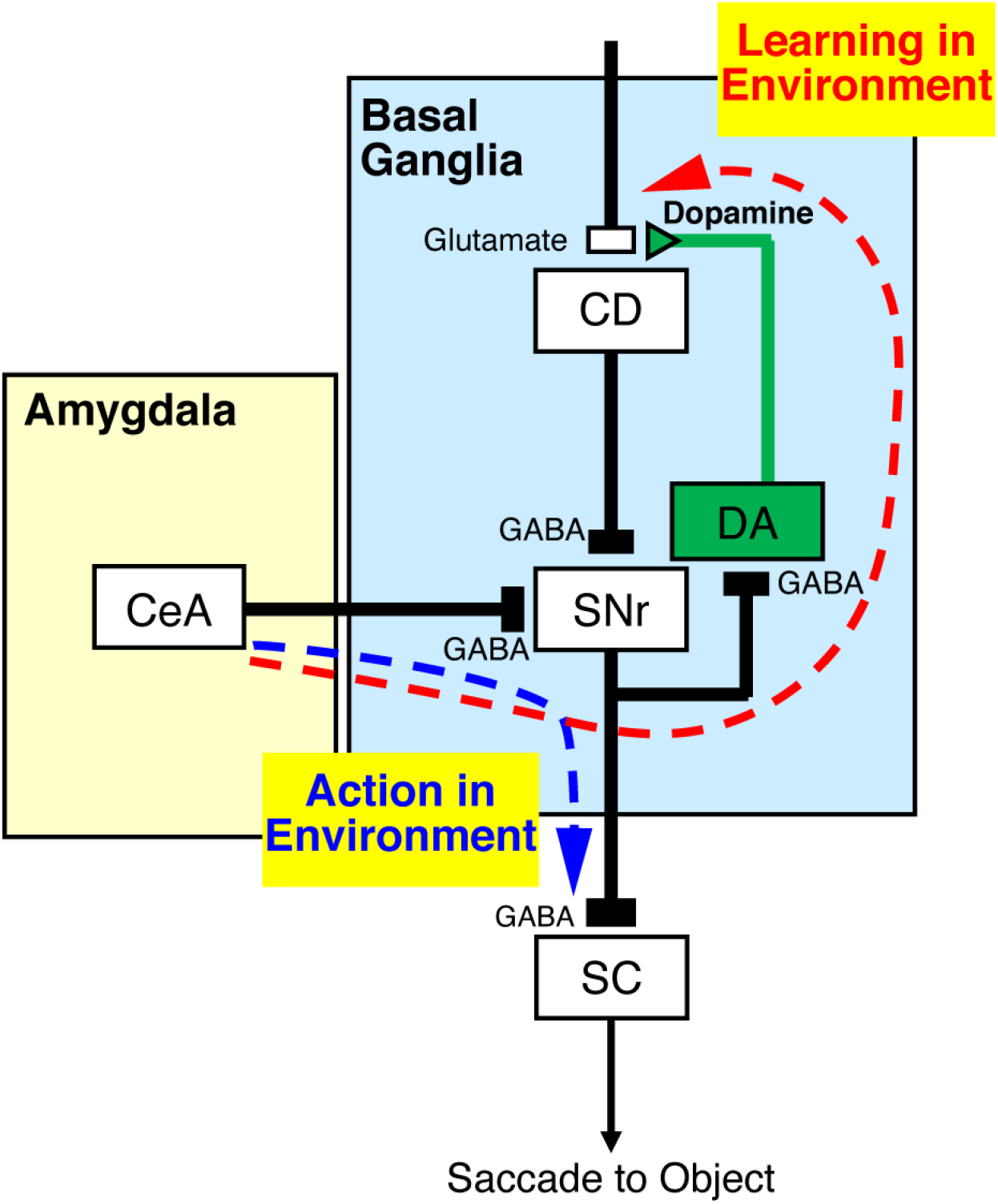
Amygdala’s two pathways for controlling BG circuits.

Such higher-order influences on behavior depend on synaptic plasticity regulated by dopaminergic projections acting on CD, which constitutes the major input structure of BG (“Learning in Environment” in Fig. 6). In this study, unlike SNr neurons that were inhibited quickly and neurons close to the site of optical stimulation, DA neurons were excited slowly and distally from the stimulation site (see Figs. 3 and 4). This suggests that amygdala neurons first inhibit SNr neurons directly, and the circuit subsequently excites DA neurons. This disinhibition is due to the fact that all SNr neurons are GABAergic (Hikosaka and Wurtz, 1985) and have local connections to DA neurons (Tepper et al., 1995; Kim et al., 2015) in addition to their projections to SC (see Fig. 6).

Most previous studies have shown that learning is caused by individual objects, primarily by the predicted outcomes (Schultz, 1998; Puig et al., 2014; Hikosaka et al., 2019b). Our present results indicate that the amygdala provides an additional source of learning to the BG circuits, namely contingencies imposed by the environment.

## Methods

### Data reporting

No statistical methods were used to predetermine sample size. The experiments were not randomized, and the investigators were not blinded to allocation during experiments and outcome assessment.

### Animals and surgical procedures

We used two rhesus monkeys (Macaca mulatta) (monkey SO: 8.5 kg, 9y old, male, monkey BA: 9.0 kg, 8y old, male). All animal care and experimental procedures were approved by the National Eye Institute Animal Care and Use Committee and complied with the Public Health Service Policy on the Humane Care and Use of Laboratory Animals. Both animals underwent surgery under general anesthesia during which a head holder and a recording chamber were implanted on the head. Based on a stereotaxic atlas (Saleem and Logothetis, 2007), we implanted a rectangular chamber targeting the amygdala. The chamber was tilted anteriorly by 15 degrees in both monkeys. After confirming the position of the recording chamber using MRI (4.7 T, Bruker), a craniotomy was performed during a second surgery.

### Electrophysiological recordings

In each session one or two multi-site (16, 24, or 32 contacts) linear electrodes (V-probe or S-probe, Plexon) were lowered into the brain using an oil-driven micromanipulator system (MO-97A, Narishige). The micromanipulators were moved independently into the amygdala and/or substantia nigra while identifying electrophysiological indicators of grey and white matter boundaries. We allowed 60 minutes for the electrodes to stabilize before starting data acquisition and the behavioral protocol. Signals were pre-amplified and stored at 40 kHz for offline processing (OmniPlex, Plexon). In real time, signals were band-pass filtered between 0.2–10 kHz, and online spike sorting was performed using a custom software implementing a voltage and time window discriminator (Blip). Analysis was based on offline spike sorting using the Kilosort algorithm followed by a manual curation in the Phy (Pachitariu et al., 2016).

### Pavlovian procedure

Each trial began with the appearance of a scene image (50 degrees diameter) that signaled either PR or NR environment and remained on present for the duration of the trial (Fig. 1). In the PR environment, one of three conditioned stimuli (fractal objects) could appear that were associated with a liquid reward (water) as an unconditioned stimulus with either 100%, 50%, or 0% probability. Water was delivered through a sipper tube positioned in front of the monkey’s mouth. In the NR environment no reward was forthcoming, and the unconditioned stimulus was an air puff (20-30 psi) delivered through a narrow tube placed 6–7 cm from the face. Three different fractal objects were used as conditioned stimuli on NR trials, and were associated with air puff probabilities of 100%, 50%, and 0%. After 1.08 s of the scene presentation, a timing cue (white square, 2 degrees) appeared in the center of the screen. After 0.68 s, one CS appeared and remained present for 1.5 s, at which point offset of CS and delivery of corresponding US occurred simultaneously. The monkeys were not required to look at these images at any point in the trial. After presented the scene image 1.08 s, the timing cue was presented. Thirty-two trials of each condition (100%, 50%, and 0% CS, crossed with PR and NR environments) were presented in pseudorandom order, for a total of 192 trials.

### Experimental control

All behavioral tasks were controlled by a custom system for neuronal recording and behavioral control system (Blip; available at http://www.robilis.com/blip/). The monkey sat in a primate chair facing a front-parallel screen in a sound-attenuated and electrically shielded room. Visual stimuli were rear-projected onto a screen by a digital light processing projector (PJD8353s, ViewSonic). Eye position was sampled at 1 kHz using a video-based eye tracker (EyeLink 1000 Plus, SR Research).

### Viral injections and optogenetics

We injected an adeno-associated virus type 2 vector (AAV2-CMV-ChR2-EYFP: 9.0 × 1012 genome copy/ml) into the amygdala of one hemisphere in both monkeys (monkey SO: left amygdala, monkey BA: right amygdala). The vector was successfully used in the macaque brain in a previous study (Inoue et al., 2015; Maeda et al., 2020), and the injection location was verified with histological procedures described previously (Maeda et al., 2020). Two penetrations in monkey BA and three penetrations in monkey SO were made into one side of amygdala at least 1.41 mm apart from each other. For each penetration, 2 μl (for monkey BA) and 2 or 3 μl (for monkey SO) of the vector were introduced at a rate of 0.4 μl/min for the first 0.2 μl, followed by 0.08 μl/min for the remainder of the injection controlled by a 10 μl Hamilton syringe and motorized infusion pump (Harvard Apparatus, Holliston, MA, USA).

To perform optical stimulation and electrophysiological recording at the same time, we used 16 or 24 contact multi-site linear electrodes with one or three combined optic fibers (S-probe, Plexon). The light source was a 473nm DPSS blue light laser with a maximum power of 100mW (Opto Engine LLC). We left the laser on continuously during the experiment and placed a mechanical shutter switch (Luminos Industries Ltd) in the light path to turn the laser on and off. We measured the light intensity at the tip of the optrode before penetration of the brain using an optical power meter (1916-C, Newport Corporation) coupled with a 818-SL/DB photo detector. The maximum light intensities were set as 0.15 mW for amygdala and 0.5 mW for the projection sites in substantia nigra. Stimulation and non-stimulation periods were pseudo-randomly interleaved 200 times during free viewing while various visual stimulation regimes were presented (including static pictures, a blank screen, and movies).

### Identification of DA, SNr, and amygdala neurons

We searched for DA and SNr neurons in and around the substantia nigra. DA neurons were identified as having less than 15 sp/s baseline activity (0–500 ms before the scene onset), phasic excitation to the scene image comparing with baseline activity, and excitation to 100% reward CS comparing with 50% or 0% reward CS. SNr neurons were identified as having more than 15 sp/s baseline activity and phasic inhibition to the scene image comparing with baseline. To reach the amygdala area, the electrode was first advanced through GPe and/or striatum areas. After passing through a quiet white matter region, spikes of amygdala neurons (heterogeneous firing patterns, relatively consistent firing with no pause) grew larger. Amygdala neurons that show phasic excitation to the scene image onset within 0–300 ms were further analyzed in this study.

Phasic visual response to the scene image was analyzed within a 0–300 ms window (DA and Amygdala) or 0-500 ms (SNr) after the scene onset and compare with baseline activity within a -500 to 0 ms window (t-test). Responses to CS or US were calculated at the time 0–300 ms after the onset. The tonic component of the scene responses was calculated in a 0–1,200 ms window before the CS onset.

### Neuronal activity analysis

For the analyses of neuronal activity, spike-density functions (SDFs) were generated by convolving spikes times with a Gaussian filter (σ = 20 ms). To investigate neuronal responses to the optical stimulation, we compared neuronal activities in a 200 ms window after stimulation vs. non-stimulation events for each neuron. Significance was assessed with an alpha threshold of 0.05. A modulation index was computed for each neuron, defined as the area under the receiver operating characteristic curve (AUROC) comparing responses to stimulation vs. non-stimulation.

Latency of responses to optical stimulation was measured by computing the earliest time that the neuron’s activity exceeded (for excitation) or fell below (for inhibition) the baseline by two standard deviations for at least 7 out of 10 consecutive ms bins. Neurons that failed to meet this criterion were excluded from analysis.

## Data availability

All data supporting the findings of this study are available from the corresponding author upon reasonable request.

## Code availability

Custom code for behavioral and electrophysiological tests is available from the corresponding author upon reasonable request.

## Acknowledgments

We thank D. McMahon, M.K. Smith, D. Parker, G. Tansey, D. Yu, A.M. Nichols, D. Yochelson, J. Fuller-Deets, A.V. Hays, and M Fujiwara for technical assistance. This work was supported by Intramural Research Program at the National Eye Institute, the National Institutes of Health, United States (project number: 1ZIAEY000415, https://reporter.nih.gov/project-details/9796699), AMED Grants (JP20dm0207077 and JP21dm0207077), and MEXT/JSPS KAKENHI Grant (JP19H05467).

## Abbreviations

BG: basal ganglia
DA: dopamine
CDh: caudate head
CDt: caudate tail
Gpe: globus pallidus externus
SC: superior colliculus
CeA: central nucleus of the amygdala
SNr: substantia nigra pars reticulata
SNc: substantia nigra pars compacta
PR: possible reward
NR: no reward

## Supplemental figures

**Fig. S1.**
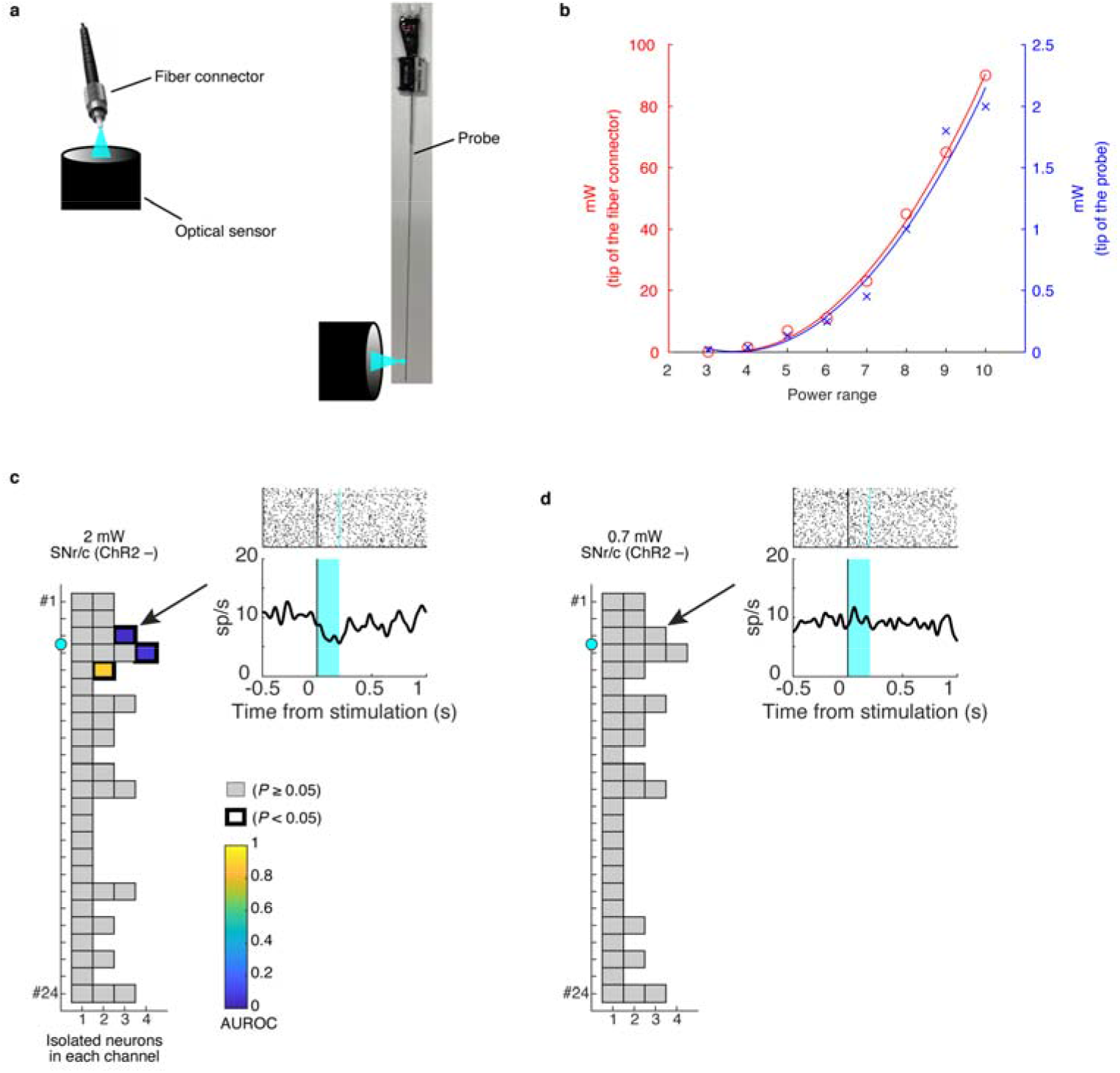
Setting of stimulation intensity (related to Figure 2) **a**, Two places for measuring optical power (left: at the tip of the fiber connector, right: at the tip of the optrode). **b**, Light intensities at the two places in different power rages. The ordinate shows measured light intensity (red: at the tip of the fiber connector, blue: at the tip of the optrode). The abscissa shows the power range on a scale of up to ten levels of the laser source. **c, d**, Responses of substantia nigra neurons during applying optical stimulation to the neurons that are not expressed ChR2 (c: 2 mW, d: 0.7 mW intensity at the tip of the optrode). The ordinate shows the place of the recording contacts from top to bottom (#1 to #24). The abscissa shows the number of isolated neurons in each contact. The place of the stimulation port of fiber optic is shown by the cyan dot (between #3 and #4 contacts). The color of each square shows the degree of modulation by the stimulation in each isolated neuron. The modulation score is defined as the area under the receiver operating characteristic (AUROC) based on the activity in stimulation trials vs. no-stimulation trials for each neuron. The scores were converted to the color scale if they were statistically modulated by stimulation (P < 0.05, T-test). The gray color indicates neurons that have no statistical significance. Example neurons were selected (black arrow) and the activities and raster plots are shown in the right panel. The cyan color in the panel shows the duration of the stimulation (200 ms).

**Fig. S2.**
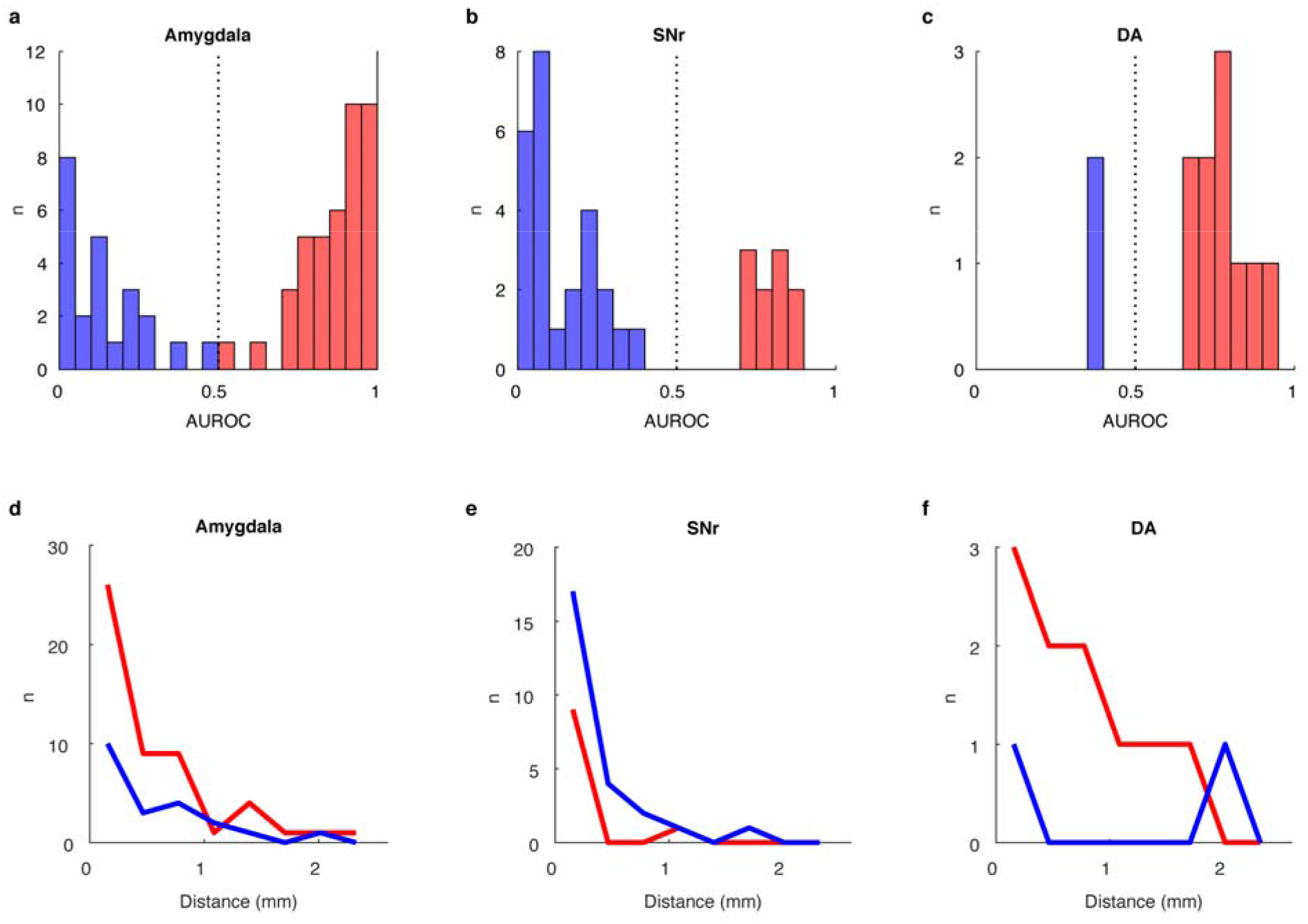
Neuronal modulation and the effective range of the optogenetic stimulation (related to Figure 2) **a-c**, The number of neurons that were modulated by the optogenetic stimulation (P < 0.05 t-test). AUROC >0.5 indicates the neuron that was excited by the stimulation and <0.5 indicates the neuron that was inhibited by the stimulation. **d-f**, The distance of modulated neurons from the stimulation site.

**Fig. S3.**
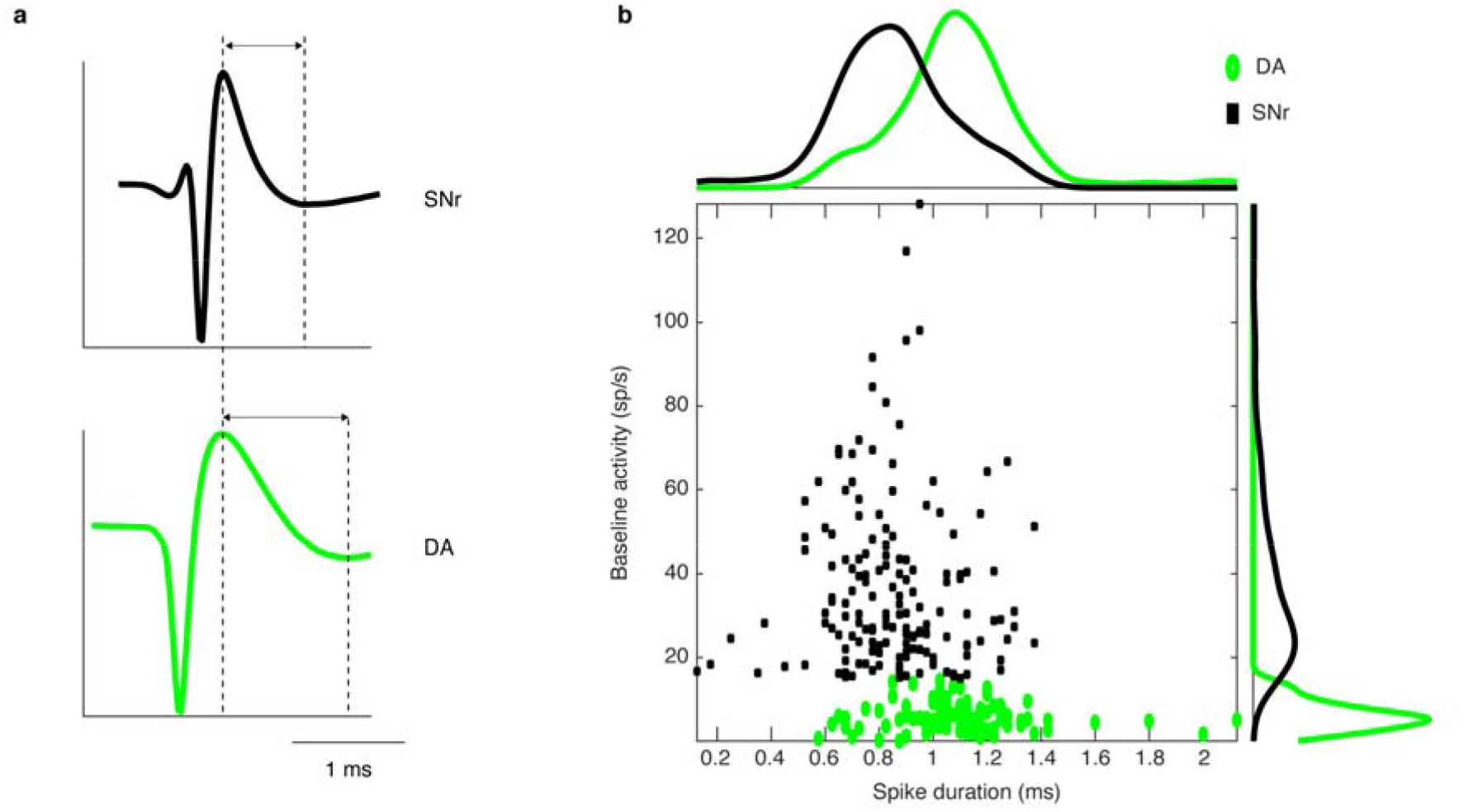
Characteristic of SNr and DA neurons. **a**, Averaged spike shapes in each group. b, The baseline firing rate (ordinate) and the spike duration (abscissa) for SNr and DA neurons. The spike duration was defined as the interval between voltage peak and subsequent trough (dotted line in a).

**Fig. S4.**
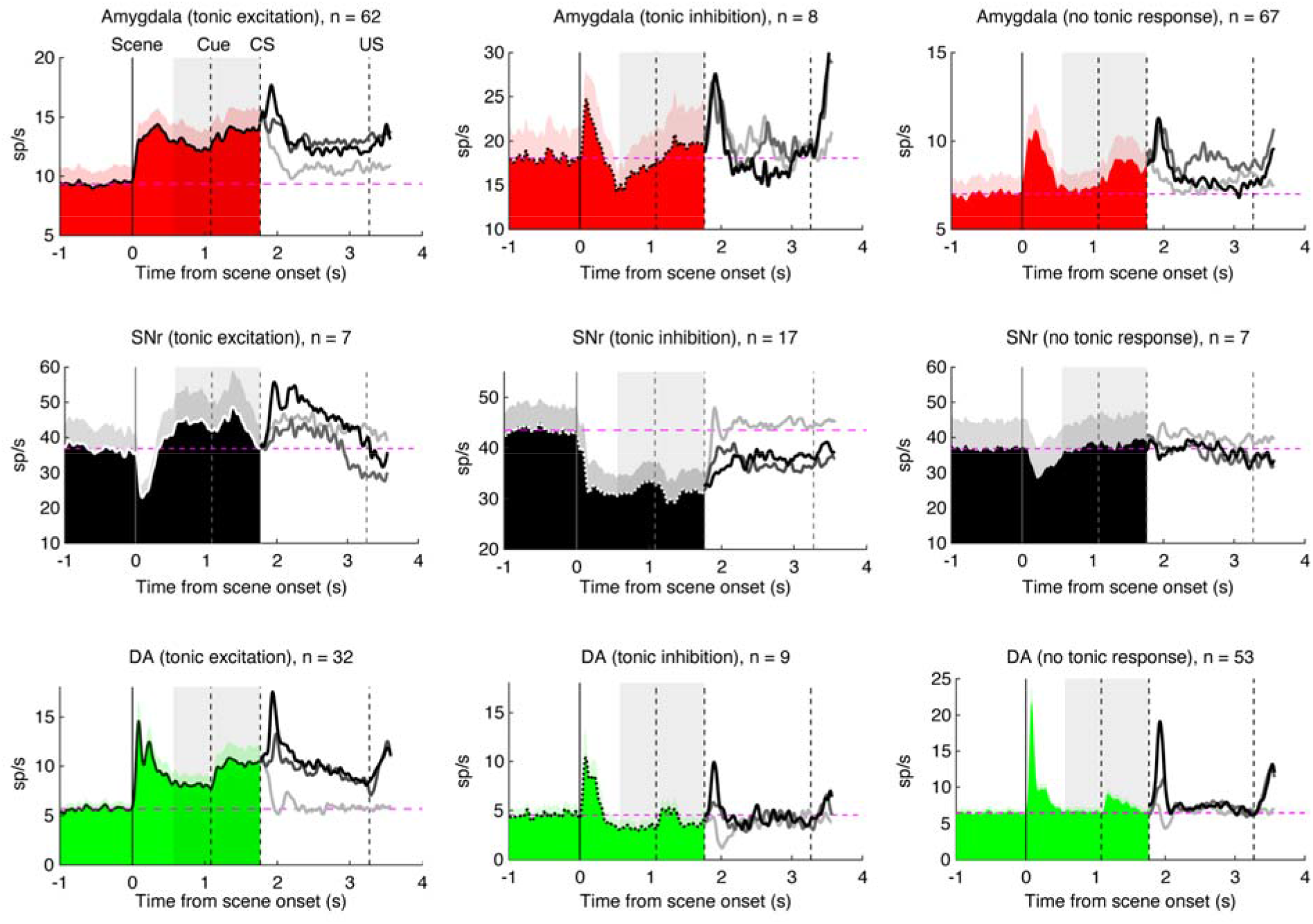
All types of responses for environmental contexts (related to Figure 4) The tonic excitation, tonic inhibition, and no tonic activity in the amygdala, SNr, and DA are shown. The format is the same as Figure 4a which shows the main responses during the task.

**Fig. S5.**
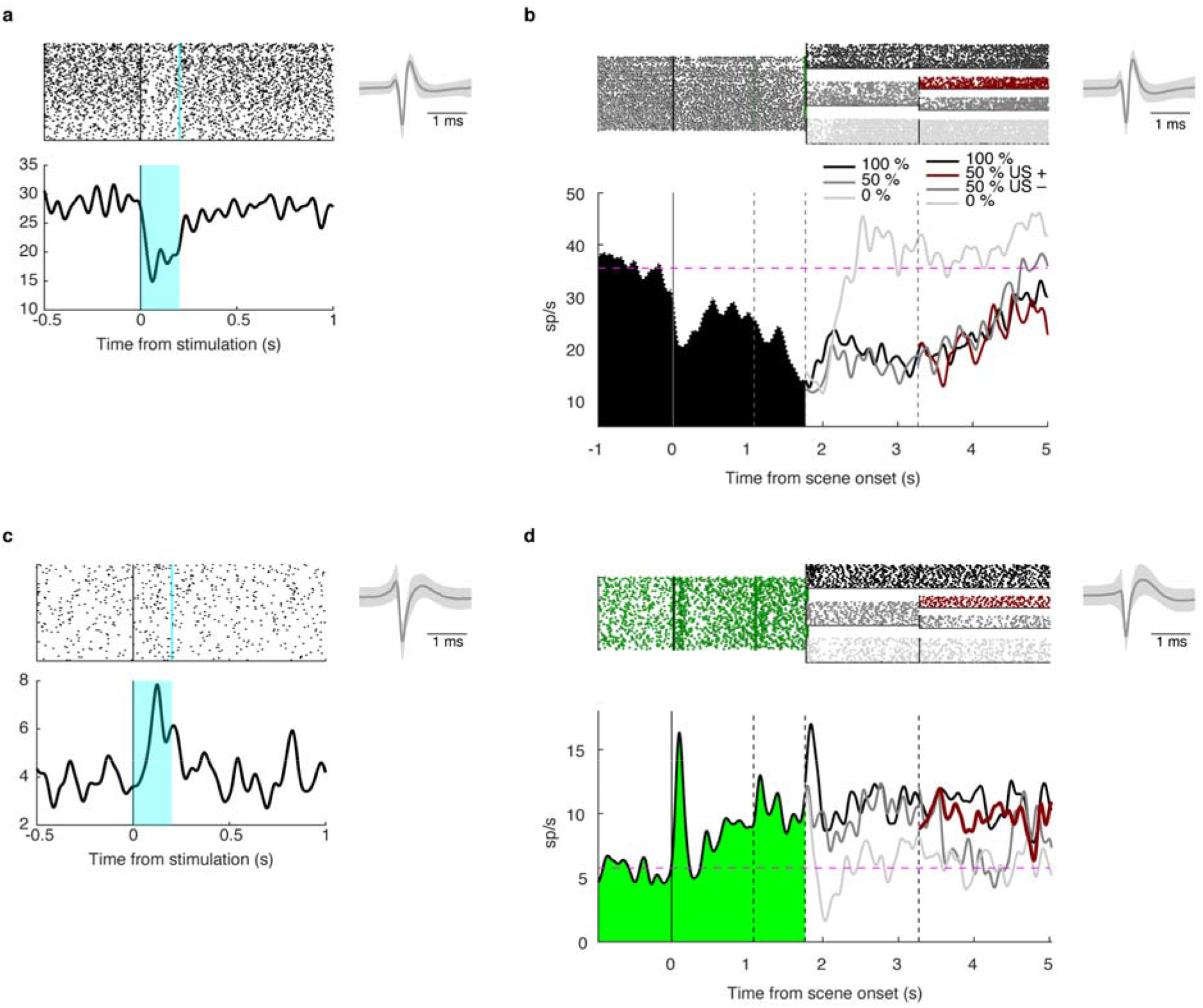
Example neuronal responses (related to figure 2 and 3) **a**, An example SNr neuron that was inhibited by optogenetic stimulation of the amygdala’s axon. **b**, Responses to PR environment in the same SNr neuron. The format is the same as Figure 1. **c**, An example DA neuron that was excited by optogenetic stimulation of the amygdala’s axon. d, Responses to PR environment in the same DA neuron. The format is the same as Figure 1. The right panel in each figure shows averaged spike shape during each task. The shaded gray area indicates ± 1 SD.

**Fig. S6.**
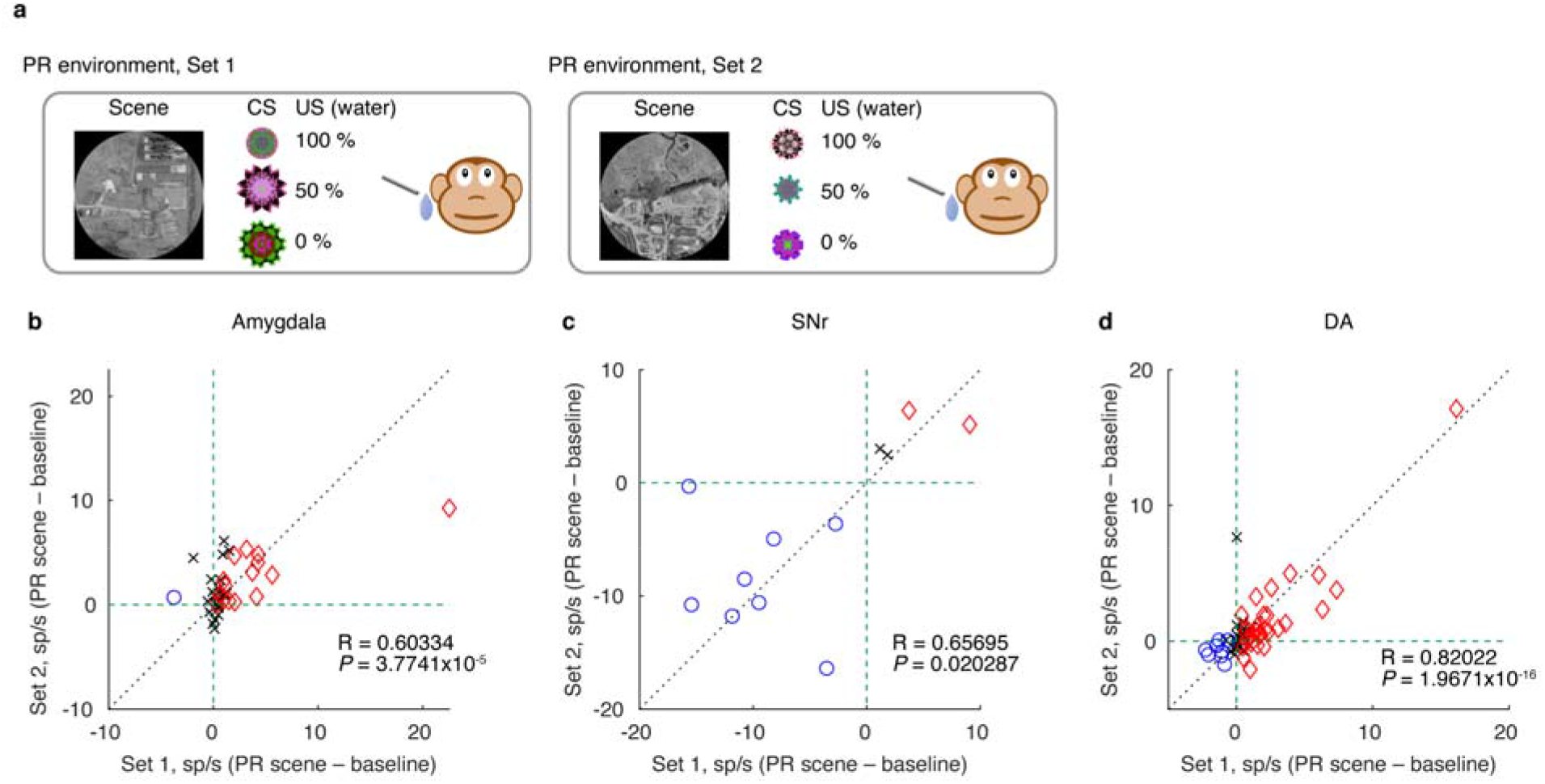
Responses to different scene images. **a**, Replication of Pavlovian task with a second set of PR scene and fractal images. **b**, Distribution of neuronal responses that were tested by both stimulus sets. The abscissa and ordinate show the response to the scene of set 1 or set 2, respectively, minus baseline activity. The red, blue, or black colors indicate that the response increased, decreased, or no change respectively (P<0.05, t-test). The R and p-value are from Pearson’s correlation.

## Notes

### Competing Interest Statement

The authors have declared no competing interest.

## References

Amir A, Lee SC, Headley DB, Herzallah MM, Pare D (2015) Amygdala signaling during foraging in a hazardous environment. J Neurosci 35:12994–13005.

Amita H, Hikosaka O (2019) Indirect pathway from caudate tail mediates rejection of bad objects in periphery. Sci Adv 5:eaaw9297.

Amita H, Kim HF, Smith MK, Gopal A, Hikosaka O (2019) Neuronal connections of direct and indirect pathways for stable value memory in caudal basal ganglia. Eur J Neurosci 49:712–725.

Bauer S, Hoye BJ (2014) Migratory animals couple biodiversity and ecosystem functioning worldwide. Science 344:1242552.

Bromberg-Martin ES, Matsumoto M, Hikosaka O (2010) Distinct tonic and phasic anticipatory activity in lateral habenula and dopamine neurons. Neuron 67:144–155.

Choi JS, Kim JJ (2010) Amygdala regulates risk of predation in rats foraging in a dynamic fear environment. Proc Natl Acad Sci U S A 107:21773–21777.

Collins AL, Greenfield VY, Bye JK, Linker KE, Wang AS, Wassum KM (2016) Dynamic mesolimbic dopamine signaling during action sequence learning and expectation violation. Sci Rep 6:20231.

Everitt BJ, Morris KA, O’Brien A, Robbins TW (1991) The basolateral amygdala-ventral striatal system and conditioned place preference: further evidence of limbic-striatal interactions underlying reward-related processes. Neuroscience 42:1–18.

Fiorillo CD, Tobler PN, Schultz W (2003) Discrete coding of reward probability and uncertainty by dopamine neurons. Science 299:1898–1902.

Fudge JL, Haber SN (2000) The central nucleus of the amygdala projection to dopamine subpopulations in primates. Neuroscience 97:479–494.

Griggs WS, Kim HF, Ghazizadeh A, Costello MG, Wall KM, Hikosaka O (2017) Flexible and stable value coding areas in caudate head and tail receive anatomically distinct cortical and subcortical inputs. Front Neuroanat 11:106.

Haber SN, Fudge JL (1997) The primate substantia nigra and VTA: integrative circuitry and function. Crit Rev Neurobiol 11:323–342.

Hamid AA, Pettibone JR, Mabrouk OS, Hetrick VL, Schmidt R, Vander Weele CM, Kennedy RT, Aragona BJ, Berke JD (2016) Mesolimbic dopamine signals the value of work. Nat Neurosci 19:117–126.

Hikosaka O, Wurtz RH (1983) Visual and oculomotor functions of monkey substantia nigra pars reticulata. IV. Relation of substantia nigra to superior colliculus. J Neurophysiol 49:1285–1301.

Hikosaka O, Wurtz RH (1985) Modification of saccadic eye movements by GABA-related substances. II. Effects of muscimol in monkey substantia nigra pars reticulata. J Neurophysiol 53:292–308.

Hikosaka O, Kim HF, Amita H, Yasuda M, Isoda M, Tachibana Y, Yoshida A (2019a) Direct and indirect pathways for choosing objects and actions. Eur J Neurosci 49:637–645.

Hikosaka O, Yasuda M, Nakamura K, Isoda M, Kim HF, Terao Y, Amita H, Maeda K (2019b) Multiple neuronal circuits for variable object-action choices based on short-and long-term memories. P Natl Acad Sci USA 116:26313–26320.

Howe MW, Tierney PL, Sandberg SG, Phillips PE, Graybiel AM (2013) Prolonged dopamine signalling in striatum signals proximity and value of distant rewards. Nature 500:575–579.

Inoue KI, Takada M, Matsumoto M (2015) Neuronal and behavioural modulations by pathway-selective optogenetic stimulation of the primate oculomotor system. Nat Commun 6:8378.

Kim HF, Hikosaka O (2013) Distinct basal ganglia circuits controlling behaviors guided by flexible and stable values. Neuron 79:1001–1010.

Kim HF, Ghazizadeh A, Hikosaka O (2015) Dopamine neurons encoding long-term memory of object value for habitual behavior. Cell 163:1165–1175.

Kobayashi S, Schultz W (2014) Reward contexts extend dopamine signals to unrewarded stimuli. Curr Biol 24:56–62.

Kunimatsu J, Yamamoto S, Maeda K, Hikosaka O (2021) Environment-based object values learned by local network in the striatum tail. P Natl Acad Sci USA 118:e2013623118.

Kwakkel G, de Goede CJ, van Wegen EE (2007) Impact of physical therapy for Parkinson’s disease: a critical review of the literature. Parkinsonism Relat Disord 13 Suppl 3:S478–487.

Lim I, van Wegen E, de Goede C, Deutekom M, Nieuwboer A, Willems A, Jones D, Rochester L, Kwakkel G (2005) Effects of external rhythmical cueing on gait in patients with Parkinson’s disease: a systematic review. Clin Rehabil 19:695–713.

Maeda K, Kunimatsu J, Hikosaka O (2018) Amygdala activity for the modulation of goal-directed behavior in emotional contexts. Plos Biol 16:e2005339.

Maeda K, Inoue KI, Kunimatsu J, Takada M, Hikosaka O (2020) Primate amygdalo-nigral pathway for boosting oculomotor action in motivating situations. iScience 23:101194.

Maren S, Phan KL, Liberzon I (2013) The contextual brain: implications for fear conditioning, extinction and psychopathology. Nat Rev Neurosci 14:417–428.

Matsumoto M, Hikosaka O (2009) Two types of dopamine neuron distinctly convey positive and negative motivational signals. Nature 459:837–841.

McDonald AJ (1982) Cytoarchitecture of the central amygdaloid nucleus of the rat. J Comp Neurol 208:401–418.

Murray EA (2007) The amygdala, reward and emotion. Trends Cogn Sci 11:489–497.

Nithianantharajah J, Hannan AJ (2006) Enriched environments, experience-dependent plasticity and disorders of the nervous system. Nat Rev Neurosci 7:697–709.

Oertel WH, Tappaz ML, Berod A, Mugnaini E (1982) Two-color immunohistochemistry for dopamine and GABA neurons in rat substantia nigra and zona incerta. Brain Res Bull 9:463–474.

Owen MA, Swaisgood RR, Blumstein DT (2017) Contextual influences on animal decision-making: Significance for behavior-based wildlife conservation and management. Integr Zool 12:32–48.

Pachitariu M, Steinmetz N, Kadir S, Carandini M, Kenneth D H (2016) Kilosort: realtime spike-sorting for extracellular electrophysiology with hundreds of channels. bioRxiv.

Parent A, Mackey A, Smith Y, Boucher R (1983) The output organization of the substantia nigra in primate as revealed by a retrograde double labeling method. Brain Res Bull 10:529–537.

Pitkanen A, Amaral DG (1994) The distribution of GABAergic cells, fibers, and terminals in the monkey amygdaloid complex: an immunohistochemical and in situ hybridization study. J Neurosci 14:2200–2224.

Price JL, Amaral DG (1981) An autoradiographic study of the projections of the central nucleus of the monkey amygdala. J Neurosci 1:1242–1259.

Puig MV, Antzoulatos EG, Miller EK (2014) Prefrontal dopamine in associative learning and memory. Neuroscience 282:217–229.

Riotte-Lambert L, Matthiopoulos J (2020) Environmental predictability as a cause and consequence of animal movement. Trends Ecol Evol 35:163–174.

Saez A, Rigotti M, Ostojic S, Fusi S, Salzman CD (2015) Abstract context representations in primate amygdala and prefrontal cortex. Neuron 87:869–881.

Schultz W (1998) Predictive reward signal of dopamine neurons. J Neurophysiol 80:1–27.

Schultz W (2016) Dopamine reward prediction-error signalling: a two-component response. Nat Rev Neurosci 17:183–195.

Schultz W, Dayan P, Montague PR (1997) A neural substrate of prediction and reward. Science 275:1593–1599.

Shinonaga Y, Takada M, Mizuno N (1992) Direct projections from the central amygdaloid nucleus to the globus pallidus and substantia nigra in the cat. Neuroscience 51:691–703.

Siegel S, Hinson RE, Krank MD, McCully J (1982) Heroin “overdose” death: contribution of drug-associated environmental cues. Science 216:436–437.

Sih A, Ferrari MC, Harris DJ (2011) Evolution and behavioural responses to human-induced rapid environmental change. Evol Appl 4:367–387.

Steinberg EE, Gore F, Heifets BD, Taylor MD, Norville ZC, Beier KT, Foldy C, Lerner TN, Luo L, Deisseroth K, Malenka RC (2020) Amygdala-Midbrain Connections Modulate Appetitive and Aversive Learning. Neuron 106:1026–1043 e1029.

Tamietto M, de Gelder B (2010) Neural bases of the non-conscious perception of emotional signals. Nat Rev Neurosci 11:697–709.

Tepper JM, Martin LP, Anderson DR (1995) GABAA receptor-mediated inhibition of rat substantia nigra dopaminergic neurons by pars reticulata projection neurons. J Neurosci 15:3092–3103.

Tovote P, Fadok JP, Luthi A (2015) Neuronal circuits for fear and anxiety. Nat Rev Neurosci 16:317–331.

Vankova M, Arluison M, Leviel V, Tramu G (1992) Afferent connections of the rat substantia nigra pars lateralis with special reference to peptide-containing neurons of the amygdalo-nigral pathway. J Chem Neuroanat 5:39–50.

Wang Y, Toyoshima O, Kunimatsu J, Yamada H, Matsumoto M (2021) Tonic firing mode of midbrain dopamine neurons continuously tracks reward values changing moment-by-moment. Elife 10:e63166.

